# Artificial Antigen Presenting Cells for Detection and Desensitisation of Auto-reactive T cells Associated with Type 1 Diabetes

**DOI:** 10.1101/2020.12.26.424333

**Authors:** Arbel Artzy-Schnirman, Enas Abu-Shah, Rona Chandrawati, Efrat Altman, Norkhairin Yusuf, Shih-Ting Wang, Jose Ramos, Catherine S. Hansel, Maya Haus-Cohen, Rony Dahan, Sefina Arif, Michael L. Dustin, Mark Peakman, Yoram Reiter, Molly M. Stevens

**Affiliations:** Department of Materials, Department of Bioengineering and Institute for Biomedical Engineering, Imperial College London, Prince Consort Road, London SW7 2AZ, UK; Kennedy Institute of Rheumatology, Nuffield Department of Orthopaedics, Rheumatology and Musculoskeletal Sciences, University of Oxford, Oxford OX3 7FY, UK; Sir William Dunn School of Pathology, University of Oxford, Oxford OX1 3RE, UK; Laboratory of Molecular Immunology, Faculty of Biology and Technion Integrated Cancer Center, Technion-Israel Institute of Technology, Haifa, Israel; Department of Immunobiology, Guy’s, King’s & St Thomas’ School of Medicine, 2nd Floor, New Guy’s House, Guy’s Hospital, London SE1 9RT, UK; Department of Immunology, Weizmann Institute of Science, Rehovot, Israel

## Abstract

Autoimmune diseases and in particular type 1 diabetes rely heavily on treatments that target the symptoms rather than prevent the underlying disease. One of the barriers to better therapeutic strategies is the inability to detect and efficiently target rare autoreactive T-cell populations that are major drivers of these conditions. Here, we develop a unique artificial antigen presenting cell (aAPC) system from biocompatible polymer particles that allows specific encapsulation of bioactive ingredients. Using our aAPC we demonstrate that we are able to detect rare autoreactive CD4 populations in human patients and using mouse models we demonstrate that our particles are able to induce desensitization in the autoreactive population. This system provides a promising tool that can be used in the prevention of autoimmunity before disease onset.

## Introduction

Type 1 diabetes (T1D) is one of the major health challenges of the twenty-first century^1^. Despite vast research conducted and the knowledge gained regarding T1D, there is no practical knowledge about prevention and the detailed pathogenesis is not yet fully understood. T1D symptoms appear when the β-cells in the pancreas are so severely damaged that they can no longer produce insulin. Destruction of the β-cells is usually a gradual process, which lasts several years. T1D progress depends both on genetic background and on environmental conditions ^2^. CD4 T-cells play a central role in the initiation and progression of the disease, as exemplified by the observation that major histocompatibility complex (MHC) II cluster variants are the most associated risk factor with T1D ^3^, and other disease risks related to the function of CD4. In T1D auto-antibodies targeted against the β-cells appear in the blood up to several years prior to onset of the disease. During this timeframe, an early diagnosis of the disease is possible by tracking these antibodies together with well-known genetic markers. So far only the symptoms are being treated, not the disease itself, and currently there is no safe and effective treatment that is approved by the Food and Drug Administration (FDA). One promising approach in the development of therapies for T1D and other autoimmune diseases is to target autoreactive T-cells and to generate specific immune tolerance while keeping the ability to respond to pathogenic antigens ^4^. Here we propose to use artificial antigen presenting cells (aAPCs) as a novel immunotherapeutic approach to manipulate and detect rare autoreactive T-cells within the precious time window before onset of the disease, with the ultimate goal being prevention of the destruction of β-cells. We report the development of unique multifunctional particle carriers presenting MHC II loaded with the GAD_555-567_ peptide, an efficient naturally processed immunodominant T-cell epitope from Glutamate decarboxylase in patients with T1D. Our aAPCs have a hollow interior that can be developed to encapsulate biologically active materials to further manipulate the immune responses and target different cell populations. The proposed aAPCs benefit from high avidity presentation of the peptide-MHC (pMHC), which allows the detection of previously undetectable autoreactive T-cell populations in T1D patients. We also demonstrate that the particles are able to reach the spleen *in vivo* and that they are able to induce desensitization of autoreactive T-cells in a specific manner *ex vivo*. Our targeted applications are 1) early diagnosis in patients with active T1D, and 2) to tolerize autoreactive T-cells before β-cells are destroyed.

## Results

With the aim of abrogating the immune-mediated destruction of β-cells, preserving residual β-cell function in recent-onset T1D and preventing the onset of the disease and islet-specific autoimmunity in at-risk or pre-diabetes patient populations, we have set to develop aAPCs based on polymer carrier particles fabricated using layer-by-layer (LbL) techniques ^5–7^ with sequential deposition of interacting polymers onto sacrificial templates. This technique is a facile and low-cost method for forming multilayer polymer films and allows encapsulation of a variety of active (bio)molecules. This technique facilitates control over the size, shape, composition, and permeability of the polymer membrane to allow controlled release of encapsulated molecules and surface molecule presentation. Hollow polymer particles made of these polymer pairs have previously been developed to encapsulate DNA ^8,9^, chemotherapeutic drugs ^10–12^, enzymes ^13–15^, and peptide vaccines ^16,17^, and used to stimulate T-cells for vaccination *in vitro* and *in vivo* ^18,19^. LbL-assembled particles therefore outperform competing technologies by the simplicity of their assembly, exquisite control over their properties, and demonstrated potential in the area of therapeutic delivery ^10,12,20^. In this study, the polymer building blocks poly(N-vinyl pyrrolidone) (PVP) and thiol-functionalized poly(methacrylic acid) (PMASH), were chosen with stability, biodegradability, tunable crosslinking density and surface functionalization in mind (Fig. 1A). We used three different silica cores as templates for polymer multilayers, allowing us to explore various antigen loads; 875±0.089 nm porous silica, 800 nm and 2760 nm solid silica cores (see details in materials and methods). Using fluorescently labelled PMASH we assessed the increase in polymer deposition as a function of layering rounds (Fig. S1). Fluorescence microscopy images show successful sequential deposition of PVP and fluorescently labelled PMA_SH_ via hydrogen bonding interaction at pH 4 on porous silica or solid silica cores (Fig. S1). Hollow polymer particles were obtained by the dissolution of the silica cores. Upon exposure to physiological conditions pH 7.4, the hydrogen bonding between PVP and PMASH diminishes, yielding disulfide-crosslinked PMA particles, as demonstrated by the shift in the zeta potential of the particles (Fig. S2). As can be seen from the TEM images, the hollow polymer particles obtained using porous silica templates have a denser appearance (Fig. 1B, right) compared to those obtained using solid silica templates (Fig. 1B, left), due to the larger amount of polymers deposited onto porous silica cores which have a larger surface area. Following the preparation of the multi-layered polymer particles, an additional layer of PMASH was added to introduce free surface thiol groups, which can be used to covalently couple proteins. We expressed three different versions of HLA-DR4(*04:01) containing a free cysteine at the C-terminus of the alpha chain which allowed site-specific orientation to the polymer particles: DR4 complex covalently linked to the T1D associated GAD_555-567_ peptide, DR4 complex covalently linked to the flu-virus hemagglutinin epitope HA_306-318_ as a control and ‘empty’ DR4 complex which did not have any covalent peptide (Fig. S3). The peptides were covalently bound to the beta chain (Fig. S3). Using 3D Structured Illumination Microscopy (3D-SIM) we visualized the presentation of functional pMHC, using a conformation-sensitive antibody (Fig. 1C and S4, middle panel), onto fluorescent polymer particles of the various sizes (Fig. 1C and S4, left panel).

**Figure 1:**
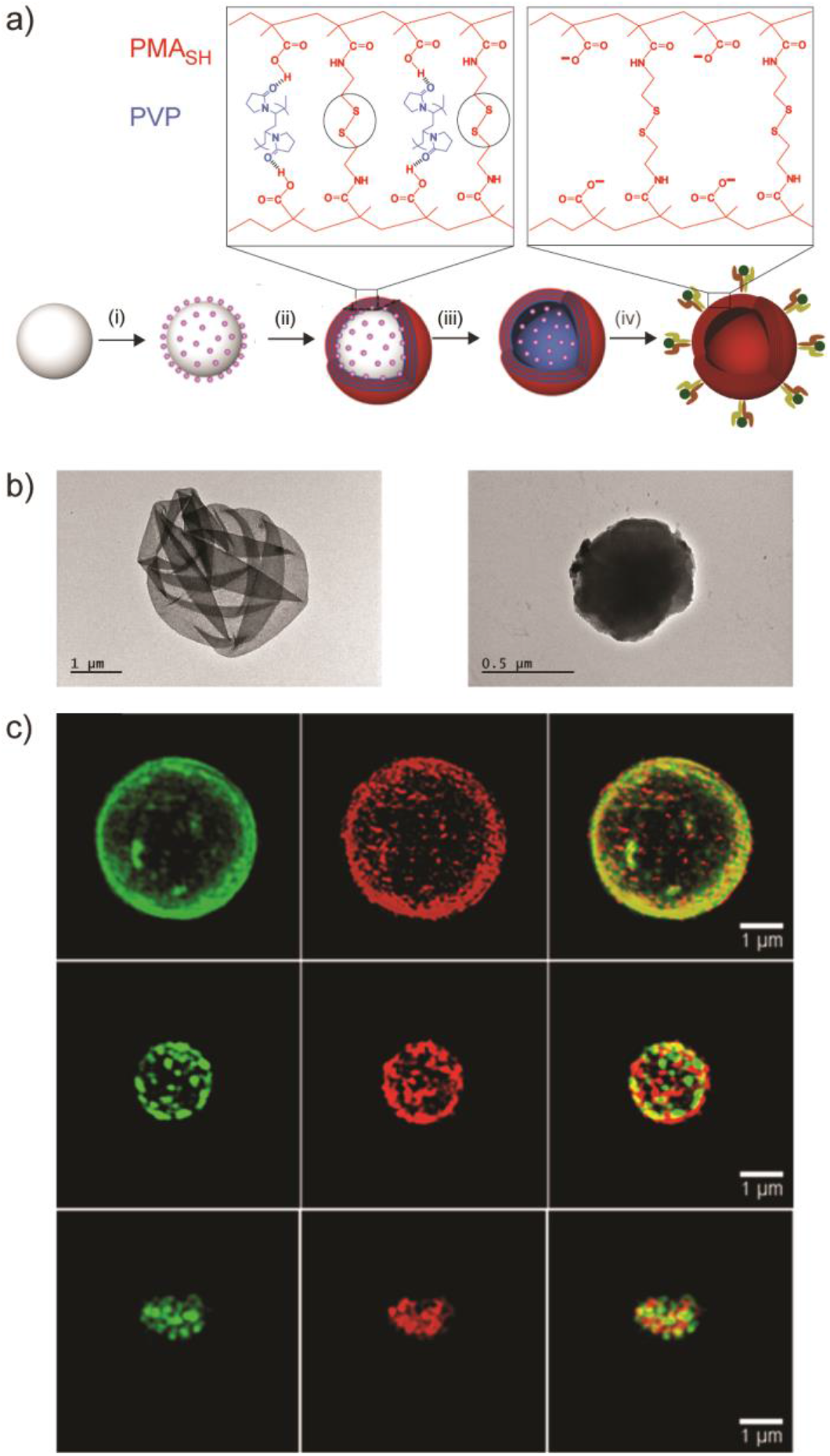
Generation of aAPC using LbL-assembled polymer particles. *(A)* Schematic illustration of the artificial antigen presenting cell preparation: (i) a sililica core is functionalised; (ii) multilayers of PMA_SH_/PVP polymers are assembled on a silica particle template; (iii) The PMA_SH_ layers are then stabilized and crosslinked through disulfide bonds followed by the removal of the silica core, resulting in a hollow polymer capsule; (iv)An additional layer of PMA_SH_ is added to allow the covalent conjugation of MHC II via disulfide linker to the capsule. *(B)* TEM images of capsules assembled using a 2.76 μm solid core silica (left) and on porous silica (right). *(C)* 3D structured illumination microscopy (3D SIM) 3D reconstructions corresponding to particles assembled on solid core silica sized 2.76 μm (top) and 1.3 μm (middle), and on porous silica core 875 nm (bottom). Green: Alexa Flour 488 labelled PMA_SH_; red: MHC II labelled with PE; yellow: overlay of the MHC II and polymers.

Next, we quantified the pMHC presentation on the various different sizes of LbL-assembled polymer particles that we generated, where we were able to create a range of presentation densities that were roughly an order of magnitude lower than the total HLA-DR4 amount on primary APCs^21^ (Fig. 2A, S5-7). The polymer particles we developed have the versatility to be directed towards different auto-immune antigens. To that end, we have tested the possibility to load exogenous peptides on our particles. We prepared polymer capsules presenting MHC refolded with endogenous low affinity peptides which were then incubated with either GAD_555-567_ or HA_306-318_ biotinylated peptides specific for the HLA-DR4(*0401), and the loading was verified using PE-labelled streptavidin (Fig. 2B). Flow cytometry (Figure 2) showed that the loading of the HA_306-318_ peptide was much more efficient compared to the GAD_555-567_ peptides. These results are consistent with the superior ability of HA-LBL to activate HA-specific clones compared to GAD-LBL activating GAD-specific clones.

**Figure 2:**
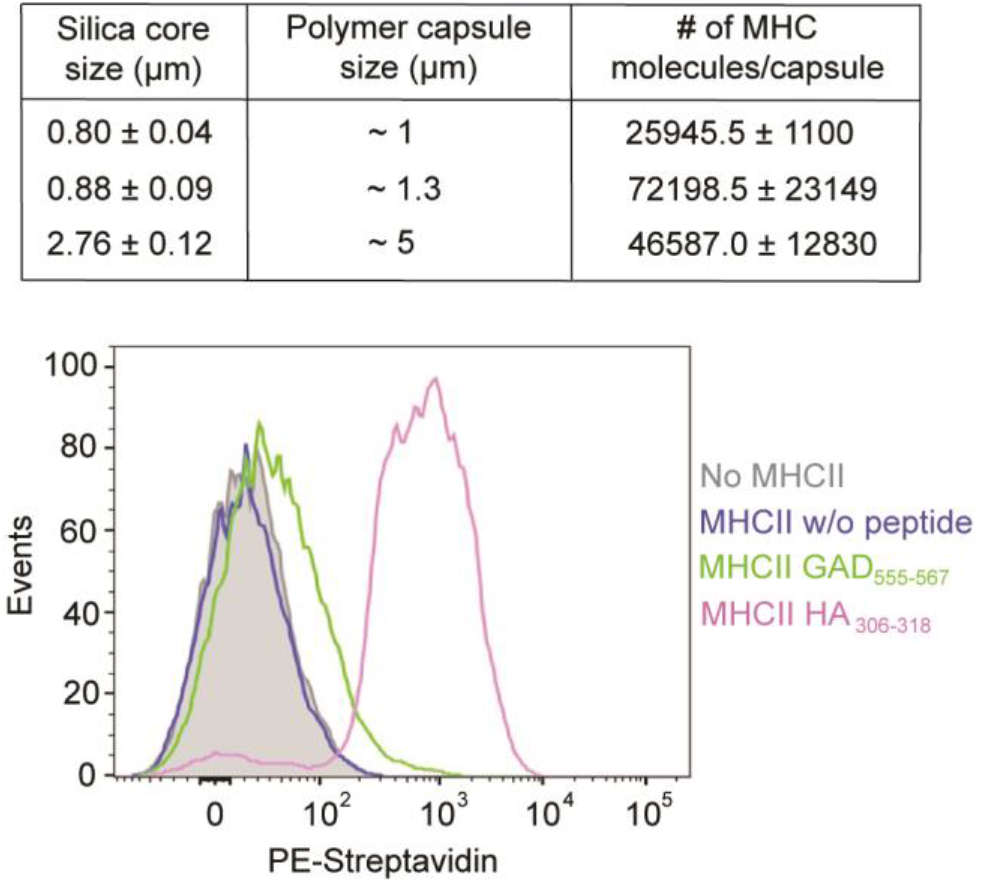
Characterisation of pMHC presentation on aAPC. *(Top)* Quantification of MHC II complexes on the capsules. Functionalized 5, 3 and 1 μm capsules were prepared and the amount of MHC II was quantified using Quantum™ Simply Cellular^®^ (QSC) microspheres. The calculated numbers of complexes are indicated. Capsules without MHC II were used as control. To determine site numbers, the fluorescence intensity of the different capsules was compared with a calibration curve of fluorescence intensity determined for beads carrying known numbers of sites. Data is shown as mean (± S.D) and is representative of at least 3 independent measurements. *(Bottom)* Flow cytometry analysis of the loaded peptide on the capsules. Capsules carrying endogenous low affinity peptides MHC II were loaded with biotinylated peptides. green-GAD65 peptide, magenta-HA peptide, blue-no peptide is loaded, Grey-capsules without MHC II. The capsules were labelled with Strep–PE. Only capsules with loaded peptides showed shift compared to capsules with no MHC II or without a peptide.

The capacity of the aAPCs particles to modulate T-cell responses was evaluated *in vitro* using human and mouse models: GAD_555-567_-specific T-cell hybridoma line G2.1.36.1 and G1.1.7.1 and HA_306-318_ specific H1.13.2 T-cell hybridoma (Fig. 3). *Ex vivo* functionality was assessed using splenocytes from DR4/RipB7 double transgenic mice. These mice are transgenic for the costimulatory molecule B7-1 driven by the rat-insulin-promoter and the DR4 allele. They exhibit age-dependent spontaneous loss of tolerance to the GAD_555-567_ (which is identical in human and mice) ^22^ (Fig. 4). We examined cytokine profiles following the incubation of the T-cell hybridomas with different ratios of polymer particles presenting either the cognate pMHC, a non-specific pMHC or unloaded (endogenous low affinity peptides). IL-2 production was only observed with aAPC presenting the cognate pMHC specifically (Fig. 3A). To further evaluate the early events leading to T-cell activation, we followed the distribution of the TCR labelled with H57 Fab’ upon interaction with the polymer particles. When aAPC lacking any pMHC are introduced, the TCR is uniformly distributed on the surface of the cells (Fig. 3B, top panel). Upon interactions with aAPC with the specific pMHC an immunological synapse is formed, polarising the TCR towards the particles (Fig. 3B, middle and bottom panels).

**Figure 3:**
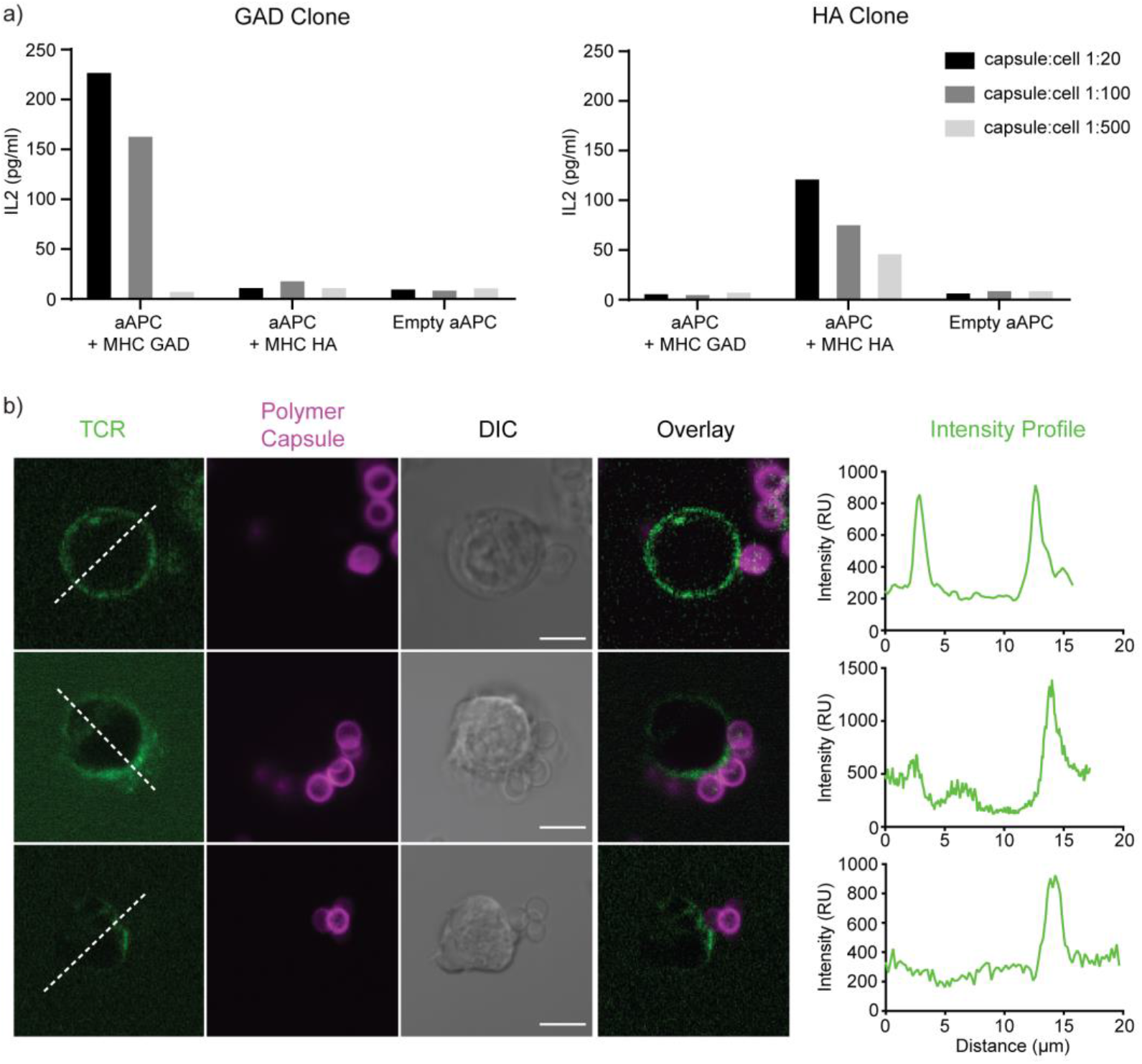
*In vitro* activation of HA or GAD-specific T-cells. *(A)* Two Hybridoma T-cells lines were incubated with aAPC and IL-2 levels were measured in the supernatants. *(B*) T-cell hybridoma polarization. The T-cell receptor (TCR) was labelled with Alexa 488-H57 Fab (green), the cells were mixed with either empty capsules (top) or aAPC (magenta) functionalized with MHC II /GAD complex (middle) or MHC II /HA complex (bottom). TCR polarization was observed in the presence of 5 μm particles carrying the DR4/GAD MHC. Cross-sections along the cell highlight the TCR polarisation towards the immunological synapse forming between the cells and the particles. Scale bars = 10 μm.

**Figure 4:**
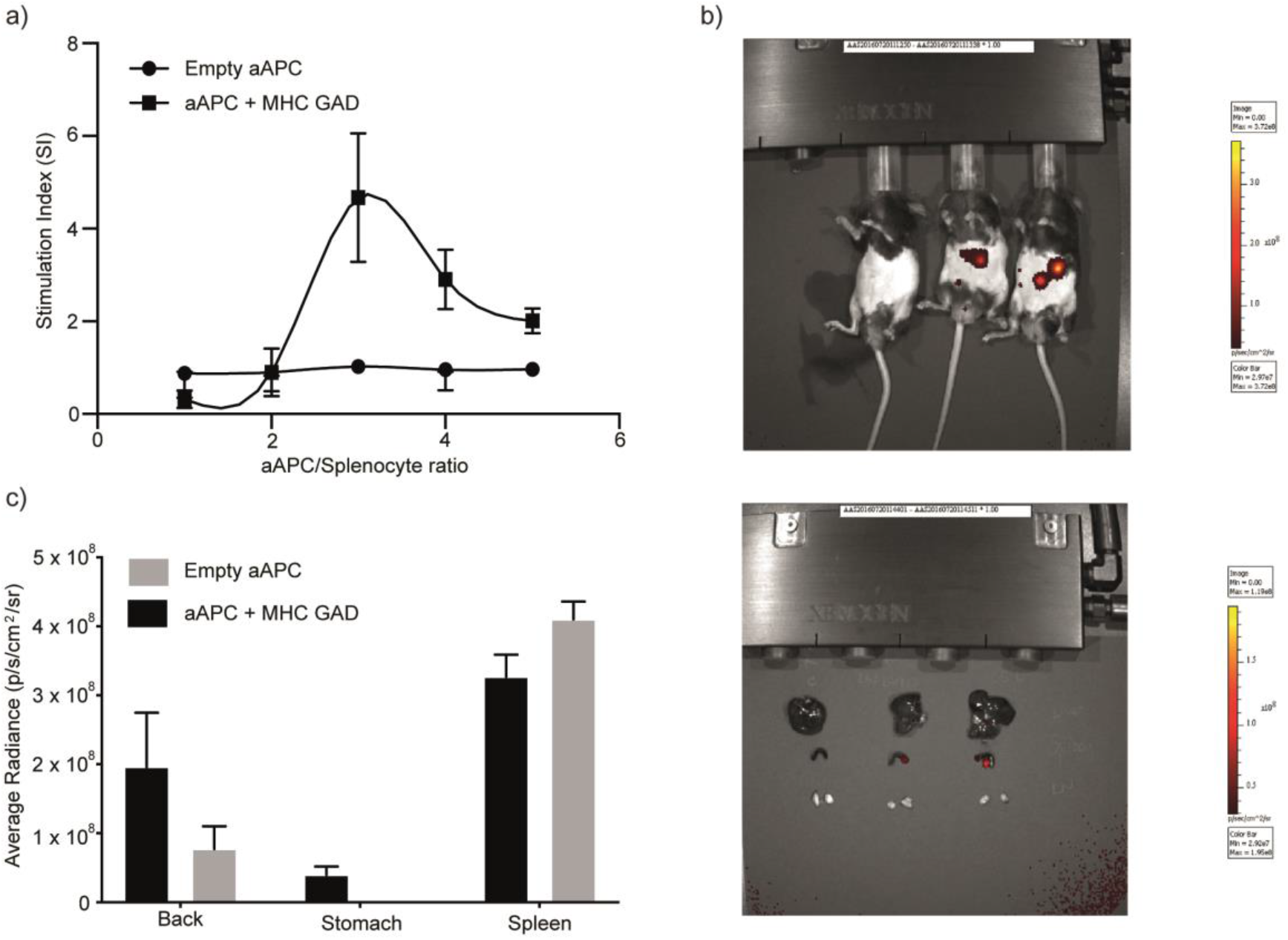
aAPC target the spleen and result in desensitisation of T-cell responses. *(A)* A recall response assay for splenocytes the aAPC functionality was performed *ex vivo* on splenocytes harvested from DR4/RipB7 (4.13 T) and from HLA-DR4 transgenic mice as a control. Cells were incubated at different cell:particle ratios as indicated, for 96 h before evaluating proliferation using 3H-thymidine incorporation. Data are shown as mean ± SD (N = 6). *(B) In vivo* bio-distribution of the aAPCs was assessed using IVIS imaging. The particles were labelled with maleimide-800. Mice were i.v injected with PBS, aAPC presenting MHC/GAD and empty particles respectively. (top) Whole body *in vivo* imaging after 24 h. (bottom) *Ex vivo* images of harvested organs (liver, spleen and lymph nodes) 24 h post injection. *(C) Ex vivo* quantification of fluorescence signals of particle accumulation in different organs. All images were obtained by PerkinElmer IVIS 200 imaging system.

To evaluate the *in vivo* activity of the engineered GAD_555-567_ specific aAPC, we examined their ability to induce tolerance using splenocytes from DR4/RipB7 transgenic mice (Fig. 4A) and followed their *in vivo* distribution in the different organs (Fig. 4B,C). To this end, splenocytes were stimulated by adding soluble GAD peptide or incubated with aAPC that either carried the GAD_555-567_ loaded MHC or the empty MHC as a control, in an increasing ratio of splenocytes/LBL. Splenocyte proliferation was measured 96 h post incubation using labeled thymidine incorporation. Using different ratios of particles to cells, we were able to reach a regime where we could inhibit the proliferation of the cells, as observed in their reduced proliferation index (Fig. 4A average of 6 mice, and S9A for each individual mice). No responses were observed towards empty particles. Using fluorescently labelled particles that were intravenously injected (i.v.), we followed the particles’ distribution 24 h post injection using In-vivo imaging system (IVIS). Particles mainly accumulated in the spleen (Fig. 4B,C and S10B,C), with no detectable accumulation in the liver or the lymph nodes, hence we can conclude that they have the potential to reach the target organ in order to induce the tolerance observed *ex vivo*.

Finally, we tested the applicability of our engineered particles not only for inducing tolerance but also for enhancing the detection of autoreactive T-cells from human samples. To that end, we used PBMCs derived from HLA-DR4^+^, recently-diagnosed T1D patients and evaluated the production of IFNg using ELISpot (Fig. 5A, S10). To evaluate the capacity of our engineered aAPC, we normalised the antigen-specific aAPC response relative to an empty aAPC and compared them to gold standard assays using peptide loaded PBMCs. It is clear that the aAPC were able to detect >50% of patients with GAD_555-567_ or HA_306-318_ responsive cells that were not detected using peptide loading (Fig. 5A, positive LbL-negative peptide square), and there was no single case where the peptide outperformed the particles. We are also able to observe tolerance induction in the patient samples by varying the ratio between the particles and the cells (Fig. 5B); however, unlike with mouse samples in which the variability was minimal (Fig. 4A), with human samples the exact ratio that induces this phenomenon is patient-specific.

**Figure 5:**
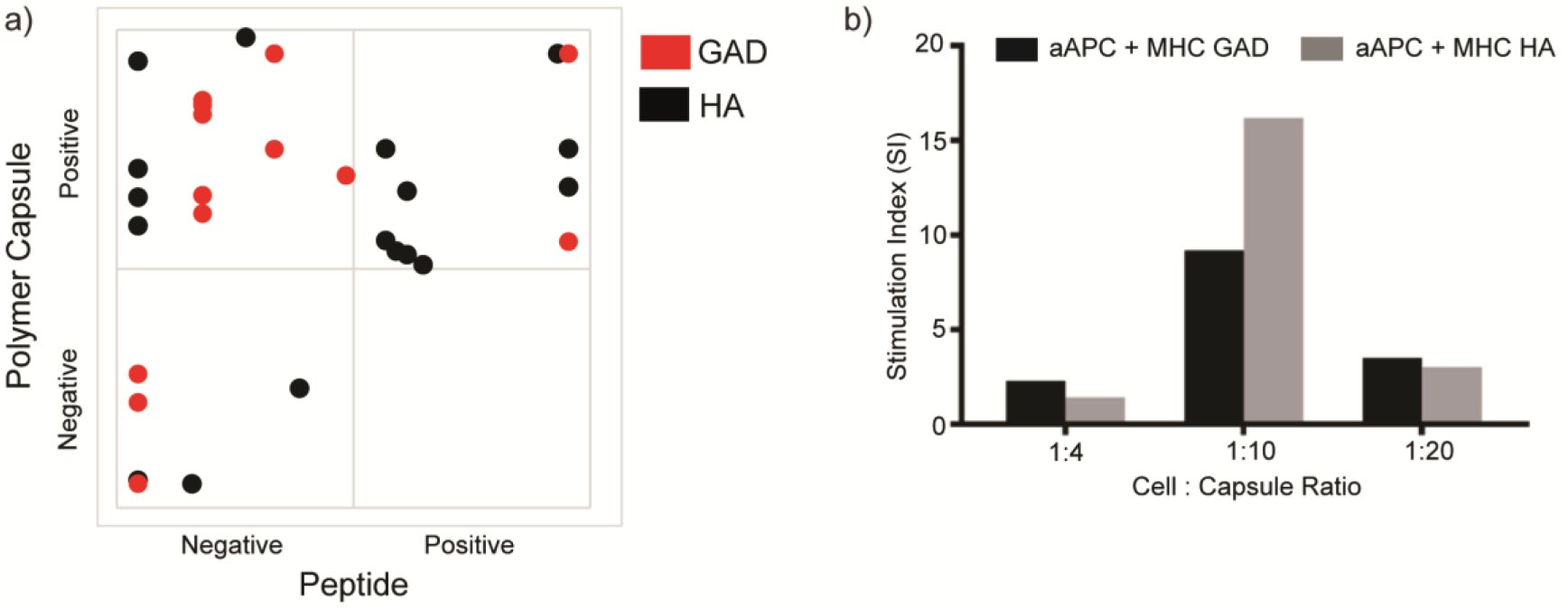
T1D patients’ response to aAPC. *(A)* IFNg positive clones as described using an ELISpot stimulation index for each patient were correlated between samples stimulated with pMHC-aAPC and the same samples stimulated using soluble peptides. Positive responders were defined as those which had a three-fold increase in response compared to background (either solvent carrier or empty LbL). *(B)* Stimulation index of two representative patients following stimulation with different aAPC to cell ratios (the exact particle to cell ratio is given in Fig. S10).

## Discussion

The need for antigen-specific immunotherapy approaches for autoimmune inflammatory diseases is well established. Strategies such as the injection of high doses of soluble peptides, NPs for the manipulation of CD8+ T-cells, and soluble MHC tetramers ^23^, have already been shown to be effective in animal models, yet translation into viable treatments in humans has had only limited success. Some of the major drawbacks of the existing techniques are their limited avidity and restricted specificity. Here we report the development of a novel immunotherapeutic tool for detection of and tolerance induction in auto-reactive T-cells, using T1D as a model system. We have designed biocompatible and versatile artificial APCs using layer-by-layer technique which allows coupling of various proteins to their surface as well as the potential to encapsulate bioactive materials such as drugs and immunomodulatory cytokines. Using our aAPCs we were able to manipulate responses of T-cells using cell lines, as well as primary cells from mouse and patient samples. Furthermore, our aAPC polymer capsules benefit from a large surface area, overcoming a key challenge in previous therapies, which have offered minimal control over the dose and avidity of the stimulus. This is demonstrated by our ability to easily detect rare populations of autoreactive cells which could not be detected using peptide pulsed PBMCs, also overcoming any dependence of specific loading efficiency of peptides of various sequences. In this work we have used pMHC-coupled polymer capsules presenting an autoantigen associated with type 1 diabetes as well as a viral antigen from influenza as a proof-of-concept. However, the versatility offered by various possible chemical modifications to the polymers comprising the Lbl-assembled polymer particles make them an attractive tool to combine different stimulations such as antibodies, multiple pMHCs, cytokines, chemokines and costimulatory or coinhibitory molecules to fine-tune the T-cell responses, taking into account the stage of the disease as well as the patients’ needs. Our work is unique in combining the ability to modulate and detect auto-reactive T-cells using a relatively simple yet resourceful system.

## Methods

### Formation of hollow polymer particles

Hollow polymer particles assembled by the layer-by-layer technique were prepared as previously described ^24^. Briefly, silica particles (50 mg/ml) were first washed with three centrifugation/redispersion cycles (1000 g, 30 s) with 20 mM NaOAc buffer pH 4.0. Assembly of polymer multilayers was achieved by alternately incubating the particles with PVP (1 mg/ml in 20 mM NaOAc buffer pH 4.0, 15 min) and PMA_SH_ (1 mg/ml in 20 mM NaOAc buffer pH 4.0, 15 min). The particles were washed three times (1000 g, 30 s) with NaOAc buffer between layers and the process was repeated until five bilayers of PMA_SH_/PVP were assembled. The thiols within the polymer layers were crosslinked with 1,8-bis(maleimido)diethylene glycol ((BM)PEG2, 1 mM in 50 mM MES buffer pH 6.0, overnight). Hollow polymer shells were obtained by dissolving the silica cores using a 2 M hydrofluoric acid (HF)/8 M ammonium fluoride (NH4F) solution for 2 min, followed by multiple centrifugation/washing cycles (4500 g, 3 min). *Caution! HF and NH4F are highly toxic. Extreme care should be taken when handling HF and NH4F solutions and only small quantities should be prepared*.

Preparation of Alexa Fluor 488-labelled PMA_SH_: A solution of PMA_SH_ (10 mg mL^-1^) was incubated with 100 μL of Alexa Fluor 488 maleimide (1 mg mL^-1^). The reaction was allowed to proceed overnight and excess Alexa Fluor 488 was removed through purification by centrifugal filter devices.

LbL concentration of each batch was determined using Flow Cytometry Cell Counting Beads (Thermo fisher, C36950).

### Cloning and expressing MHC class II/peptide complexes

Plasmids for the expression of recombinant four-domain MHC class II DR4 molecules in Schneider S2 cells, a gift from Dr. Lars Fugger, DR-A1*0101/DR-B1*0401, DR-A1*0101/DR-B1*0401(HA-306-318) and DR-B1*0401(GAD-555-567), have been previously described ^25^. Site-directed mutagenesis (Quick change II, Agilent 200523) was used to introduce the C-terminus cysteine mutation in the DR-A chain using the following primers:

DRA5’ cys(GAAGATCGAGTGGCACTGTTAAAAGGGCAATTCTG), DRA3’cys(CTTCTAGCTCACCGTGACAATTTTCCCGTTAAGAC). DR-A and DR-B plasmids were co-transfected with pCoBlast selection vector to S2 cells using cellfectin reagent (invitrogen). Stable single-cell line clones were verified for protein expression. Upon induction with CuSO4, cell supernatants were collected and DR4 complexes were affinity purified by anti-DR LB3.1 (ATCC number HB-298) mAb. The purified DR4 complexes were characterized by SDS-PAGE. The correct folding of the complexes was verified by recognition of anti-DR conformation sensitive mAb (L243) in an ELISA binding assay^26^.

Peptides for loading and functional assays were synthesized by standard fluorenylmethoxycarbonyl chemistry and purified to >95% by reverse phase HPLC, GAD_555-567_ (NFFRMVISNPAAT) HA_306-318_ (PKYVKQNTLKLAT) or bought from Gene Script.

### Cells lines and media

The Preiss cell line (EBV-transformed B-lymphoblast), H2-1 T-cell hybridomas, DR*1501 L cells transfectant (L466.1) and CTLL cell lines were maintained in complete media in RPMI-1640 supplemented with 10 v/v fetal calf serum (FCS), 2 mM glutamine, Penicillin (100 units/ml) and Streptomycin (100 μg/ml).

The G2.1.36.1 and H1.13.2 T-cell hybridomas were maintained in Dulbecco’s modified Eagles medium (DMEM) and with 10 v/v FCS, 2mM glutamine and 1v/v Penicillin Streptomycin antibiotics.

### Mice

All experiments were performed in accordance with the Israeli laws and approved by the ethics committee at the Technion-Israel Institute of Technology. DR0401-IE mice12-Class II deficient C57Bl/6 mice (I-Abo/o) transgenic for the DRA1*0101 and DRB1*0401 genes. These mice express a human-mouse chimeric class II molecule in which the TCR interacting and peptide binding domains of mouse I-E (domains α1 and β1, exon 2 in both genes) have been replaced with the α1 and β1 domains from DRA1*0101 and DRB1*0401, respectively. Retention of the murine α2 and β2 domains allows for the cognate murine CD4-murine MHC interaction.

RIP-B7/DR0401 - C57Bl/6 mice transgenic for the costimulatory molecule B7-1 driven by the rat-insulin-promoter (RIP-B7 mice) were crossed with DR0401-IE mice to generate RIP-B7/DR0401-IE (B7/DR0401) mice. These mice are spontaneously diabetic with disease incidence of ~30% at 54 weeks and a mean diabetes onset age of 37 ± 9 weeks. These mice exhibit age-dependent spontaneous loss of tolerance to 2 islet Ags GAD_555-567_ (identical sequence in human and mice) and GFAP-240-252 during the pre-diabetic and diabetic phases.

All mice in the *ex vivo* and *in vivo* experiments were 8 weeks when the experiments were performed.

### Flow cytometry

Quantification of pMHC on antigen presenting cells (B-cell line, or monocyte derived dendritic cells) or on LbL particles was performed using the Quantum^™^ Simply Cellular^®^ (QSC) microspheres and an anti HLA-DR4 PE-conjugated mAb (L243). Samples were analyzed on a X20 Fortessa or LSRII flow cytometer (BD Biosciences).

### IL-2 ELISA

T-cell hybridoma cells (2×10^5^/well in a 96-well plate) in 100 μl of 10% FBS-containing medium were combined with pMHC LbL at different ratios or as a control with 2×10^5^ DR4-EBV-transformed DR4 positive B lymphoblast MGAR cell line in 100 μl alone or in the presence of peptides in the indicated concentration. Cells were incubated at 37 °C and 5% CO_2_. After 24 h of culture. The presence of supernatants IL-2 was measured by sandwich Elisa. ELISA plates were coated with anti-IL-2 Ab (BioLegend) at a concentration of 0.25ug/ml overnight at 4 °C or 1 h at 37 °C. The plates were blocked for 30 min at room temperature with 5 v/v FCS-PBS and subsequently were incubated with the hybridoma supernatants for 2 h at room temperature. Then a biotinylated anti-IL-2 Ab (BioLegend) at a concentration of 0.25 μg/ml was added and after washing, plates were incubated with Strp-HRP antibody.

### Proliferation assay

Splenocytes from immunized mice were isolated and then cultured in a density of 5*10^5 cell/well in triplicates with or without peptide antigens for 4 days in a stimulation media (DMEM supplemented with 10 v/v FBS, 2 mM sodium pyruvate, 2 mM l-glutamate, 4 mM 2-mercaptoethanol, and 50 μg/ml penicillin/streptomycin) with GAD-LBL or empty-LBL at splenocytes to LBL ratio as indicated. The last 16 h of incubation were in the presence of 3H-thymidine to assess proliferation responses. Cultures were harvested on glass fiber filters and uptake of 3H-thymidine was assessed by liquid scintillation. The stimulation index (SI) was calculated by dividing the cpm of peptide-stimulated cultures by the cpm of control cultures.

### Human samples

PBMCs from 11 diagnosed patients with T1D were collected. All patients were HLA-DR4+. Fresh and frozen samples were compared with no obvious difference. Local research ethics committee approval (National Research Ethics Committee, Bromley NRES Committee, reference number 08/H0805/14) was granted at each participating center, and written informed consent was obtained in all cases.

### Cytokine ELISpot

Fresh or frozen PBMCs were dispensed into 48-well plates at a density of 2 × 106 in 0.5 ml in RPMI-1640 supplemented with antibiotics Pen/Strep (TC medium; Life Technologies Ltd.) and 10 v/v human AB serum (Sigma) supplemented with peptides (GAD, HA, and INFANRIX) or carrier solvent (DMSO or PBS respectively) to a final concentration of 10 μM or the LBL in different ratios, and incubated at 37 °C, 5% CO_2_, tilted by 5°. Control wells contained TC medium with an equivalent concentration of peptide/LBL. On day +1, 0.5 ml prewarmed TC medium/10% AB serum was added, and on day +2, nonadherent cells were resuspended using prewarmed TC medium/2% AB serum, washed, brought to a concentration of 106/300 μl, and 100 μl dispensed in triplicate into wells of 96-well ELISA plates (Nunc Maxisorp; Merck Ltd., Poole, United Kingdom) preblocked with 1w/v BSA in PBS and precoated with monoclonal anti–IFN-γ (U-Cytech, Utrecht, The Netherlands). When sufficient cells were available, IFN-γ was analyzed for each test peptide/LbL. After capture at 37°C, 5% CO_2_ for 22 hours, cells were lysed in ice-cold water, plates washed in PBS/Tween 20, and spots developed according to the manufacturer’s instructions. Plates were dried and spots of 80–120 μm counted in a BioReader 3000 (BioSys, Karben, Germany).

ELISpot analysis was done by individual blinded to the stimulation condition. A threshold of 3 clones was set up to account for a meaningful positive response. Samples stimulated with pMHC LbL were normalised to the response to LbL without MHC, and for the peptide stimulation the samples were normalized to solvent. To account for donor variability, the data was normalised to the maximum response of each donor.

### Microscopy

3D structured illumination microscopy (3D SIM) imaging was performed at room temperature with an Elyra PS.1 (Carl Zeiss). A 63 × 1.4 NA oil-immersion objective lens was used, with three orientation angles of the excitation grid and five phases acquired for each image with a 110 nm z-step and a pixel size of 64 nm imaged at 16 bits per pixel on an Andor Zyla. SIM processing was performed with the SIM module of the Zen software package (Carl Zeiss), then TIF stacks of processed SIM data were exported. The SIM data sets were then processed into projection images using ImageJ software.

Transmission electron microscopy (TEM) imaging. All imaging was performed with JEOL 2100F TEM, with an acceleration voltage of 200 kV. For sample preparation, 10 μl of the particle solutions were dropped on the 200 mesh copper grid covered with a carbon-stabilized Formvar film (Electron Microscopy Sciences). The residual solution was removed after 5 min. The sample with LbL polymer shells was negatively stained by 5 μl of 1 wt% uranyl acetate solution on the grid for 5 min.

Scanning confocal microscopy was performed using a 63x silicon oil objective on an Olympus FV1200. T-cells were labelled with H57-Alexa488 Fab for 20 min before a brief wash at 4 °C to remove unbound Fab, followed by a 10 min incubation with LbL particles before imaging.

## Acknowledgements

AAS was supported by EMBO Young Investigator Fellowship. AAS and MMS were supported by the Rosetrees Trust. EAS was supported by Oxford-UCB fellowship. MLD was supported by Wellcome PRF 100262Z/12/Z and Kennedy Trust for Rheumatology Research. EAS an MLD were partially supported by the Human Frontiers Science Program.

## Author Contributions

AAS & EAS Conceptualisation, investigation, methodology, visualisation, formal analysis, data curation, project administration, funding acquisition, writing original draft, writing - review & editing. RC: methodology, writng-review & editing. EF: Investigation, writing - review & editing. NY: methodology. STW investigation, methodology, fourmal analysis writing - review & editing. JR: methodology. CSH: visualisation, writing - review & editing. MHC& SA: methodology, writing - review & editing. RD&YR: Conceptualisation, resurces. MLD& MP: funding acquisition, supervision, writing - review & editing. MMS: Conceptualisation, funding acquisition, supervision, writing - review & editing

