## Supplementary for "Artificial Antigen Presenting Cells for Detection and Desensitisation of Auto-reactive T cells Associated with Type 1 Diabetes"

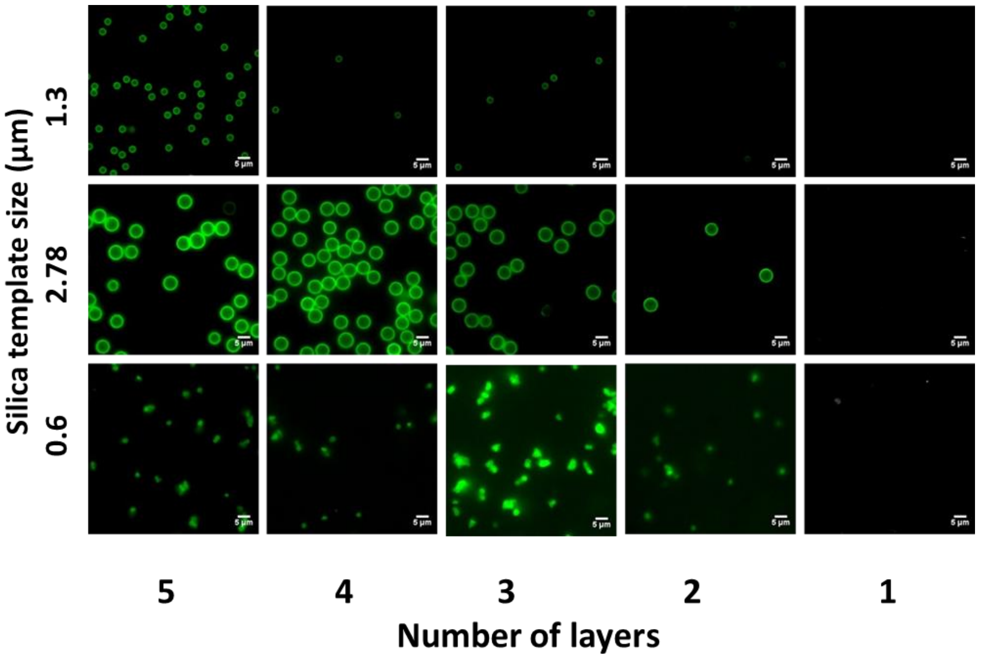


**Figure S1:** Layer-by-layer assembled polymer particles. PVP and fluorescently labelled $PMA_{SH}$ were sequentially deposited on silica particle templates and confocal images were taken after each deposition step. There is an obvious increase in fluorescence as the layering rounds progress. Note the decrease after the 3rd round for the porous silica core (0.6 µm) which is probably a result of self-quenching of the fluorophore due to its high surface density.

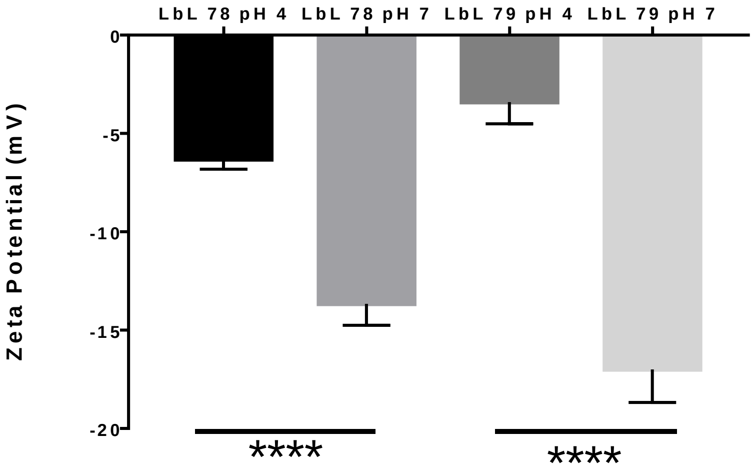


**Figure S2:** Zeta potential of LbL particles. The zeta potential was measured for two batches of LbL particles (labelled 78 and 79) at pH 4 where the particles were still compact and at pH 7 after the particles' core expanded. Data shown as mean ± S.D., n = 3 , Statistical test, **** $p < 0.0001$.

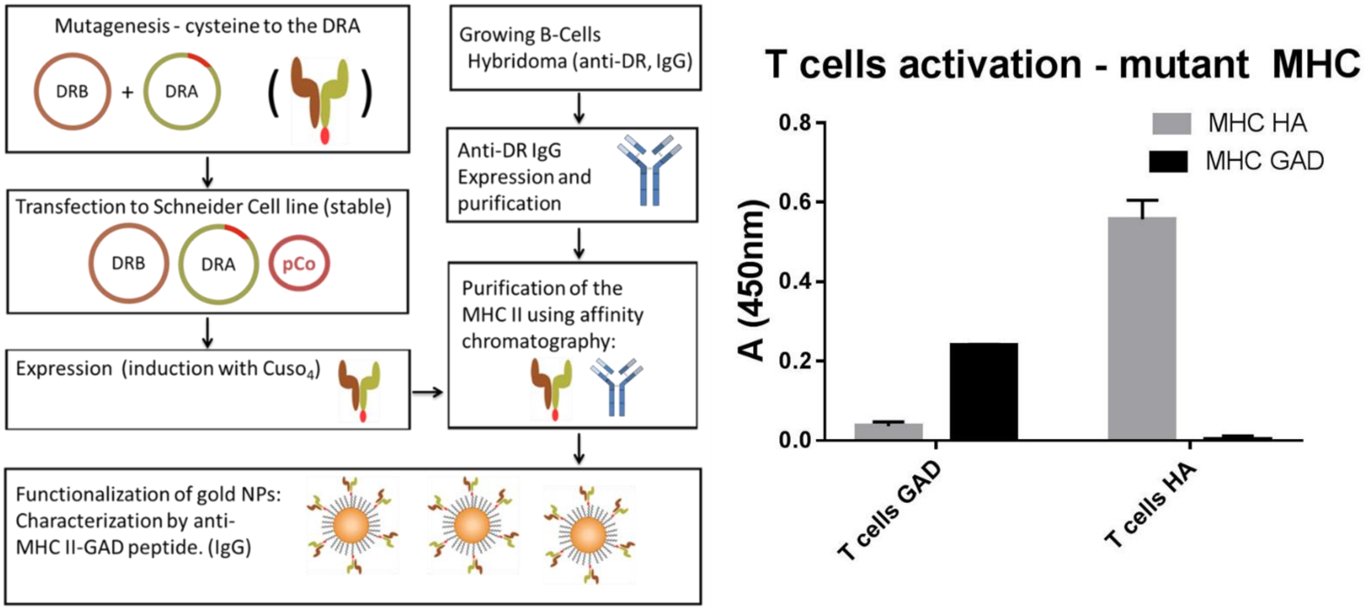


**Figure S3:** Generation of engineered HLA-DR4/peptide constructs. A) Schematic representation of the generation of cysteine modified HLA-DR4, expression in insect cells and purification of correctly refolded monomers using a confirmation sensitive antibody. The fully functional pMHC is then coupled to our particles through disulfide bonds. B) Validation that the T-cell respond well to the (cysteine) mutated MHC. Data shown as mean ± S.D., n = 3.

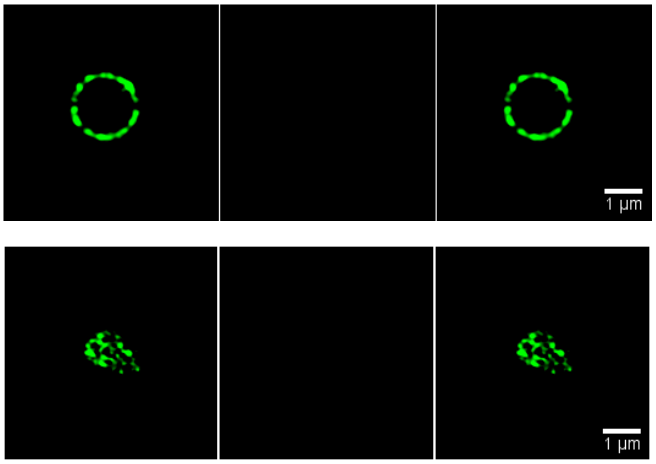


**Figure S4:** SIM images of unloaded particles. Top panel shows the fluorescent LbL on 2.7 µm core particles, bottom panel is with the porous silica. The middle panel shows the lack of MHC signal after labelling with the anti-DR4 antibody.

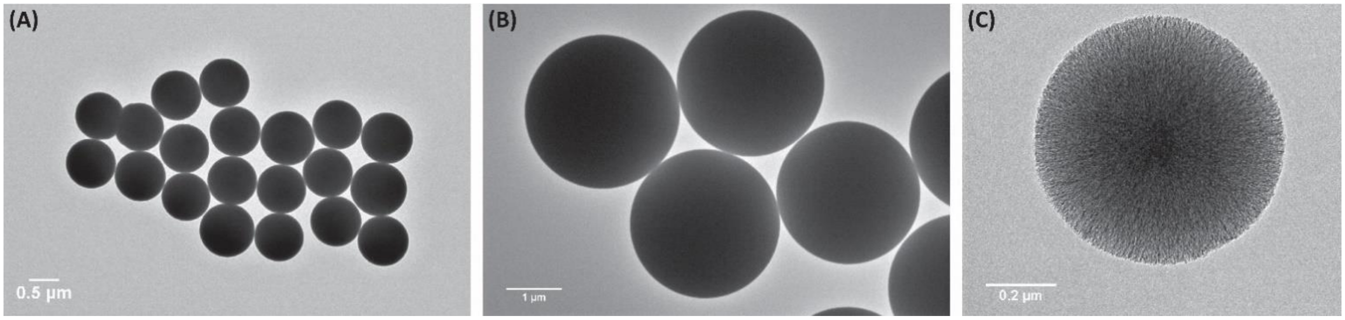

**Figure S5:** TEM images of particles of different cores size: (A) 800 nm, (B) 2700 nm and (C) porous silica.

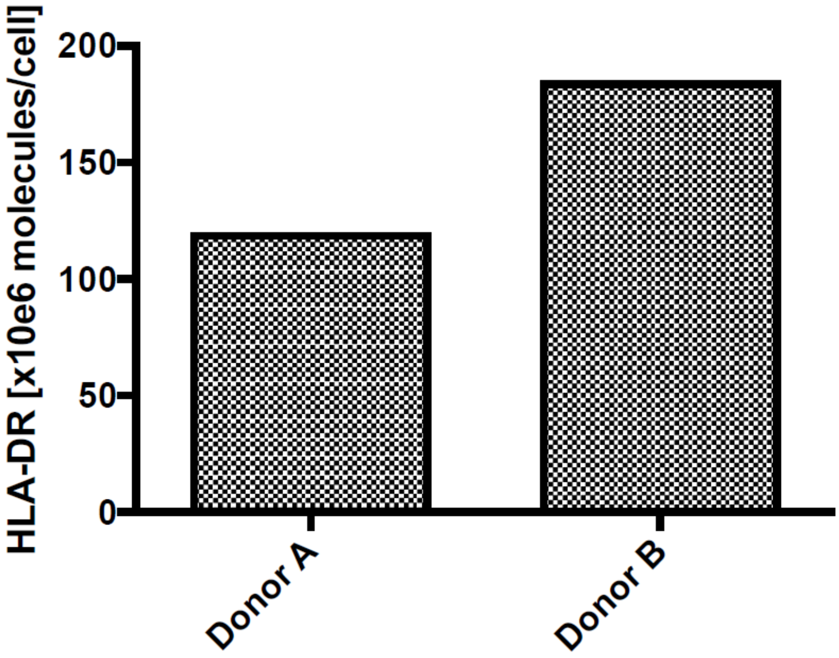


**Figure S6:** Number of HLA-DR molecules on the surface of mature monocyte derived DCs from two different donors.

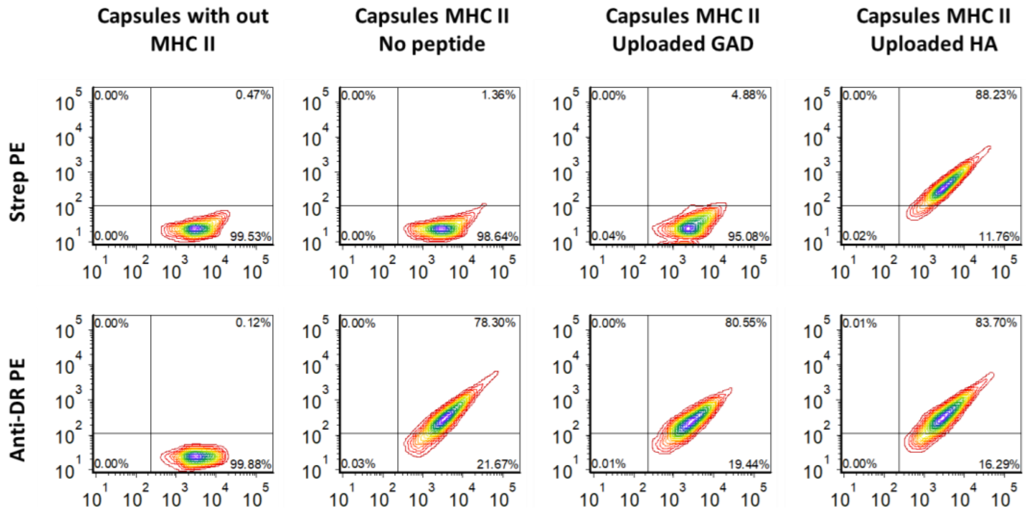


**Figure S7:** Flow cytometry histograms showing the loading of soluble peptide on LbL particles containing "unloaded" MHC. The x axis is the signal from the labelled $PMA_{SH}$ layer (Alexa Fluor 647), x axis in the top panel is streptavidin PE (binds to biotinylated peptides) and bottom panel is an anti HLA-DR4 PE specific antibody. Note the correlation between the number of layers (APC intensity) and the amount of MHC (PE intensity lower panel), also note the low loading efficiency of the GAD peptide.

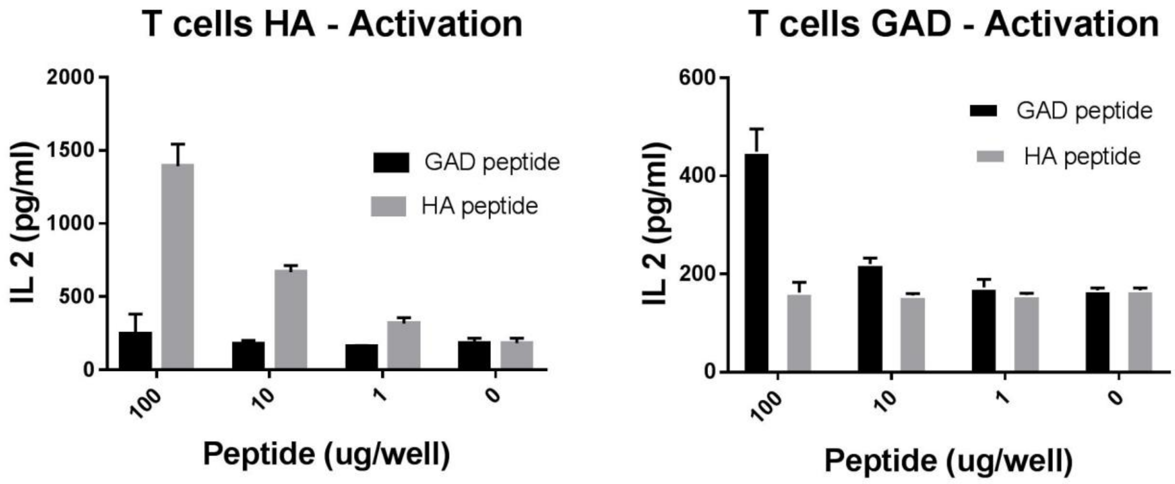


**Figure S8:** IL2 production using soluble peptides and antigen presenting cells demonstrating the lower activation of the GAD cell lines (lower IL2 production for the same amount of peptide as the HA) Data shown as mean ± S.D., n = 3.

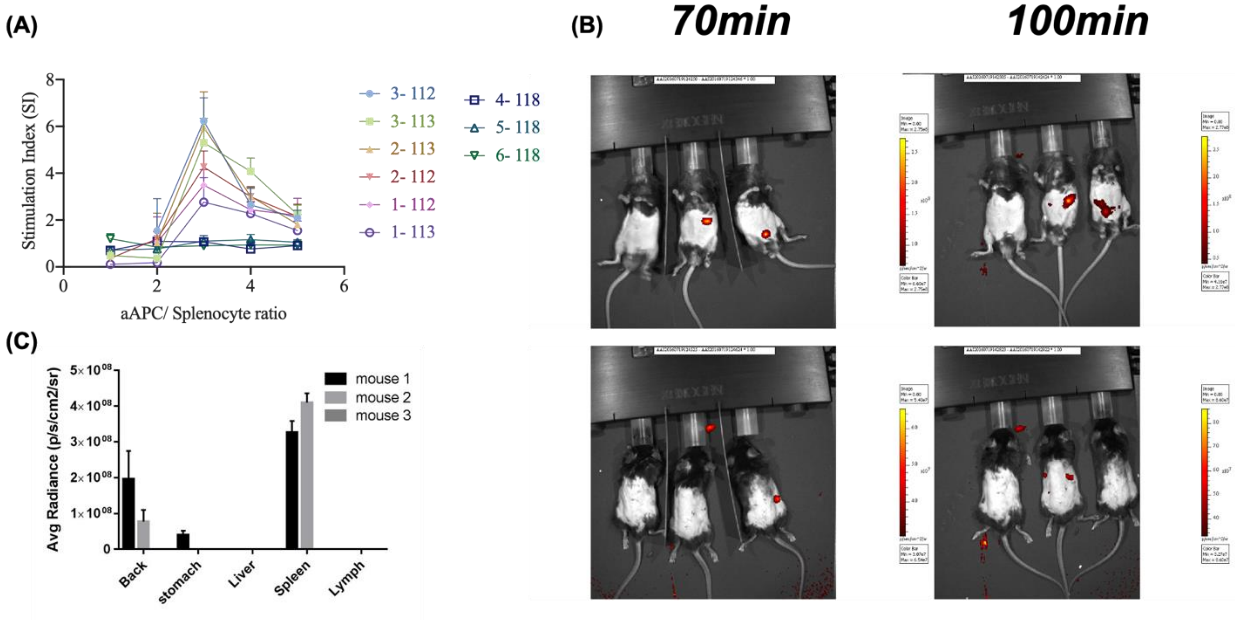


**Figure S9:** (A) A recall response assay for splenocytes the LbL particle functionality was performed ex vivo on splenocytes harvested from DR4/RipB7 (4.13 T) and from HLA-DR4 transgenic mice as a control. Cells were incubated at different cell: particle ratios as indicated, for 96 h before evaluating proliferation using 3H-thymidine incorporation. Experiments are average of 6 independent mice. (B) IVIS imaging of particle distribution 70 and 100 min after injection. The bright spot where a significant number of particles accumulates. (C) Quantification of particle accumulation in different organs (mouse 1: GAD particle; mouse 2: empty particles; mouse 3: PBS control).

| | Patient number | controls | | | | SI | |
|---|---|---|---|---|---|---|---|
| | | HA | GAD | INFANRIX | Cells:LbL | LbL HA | LbL GAD |
| 1 Background no MHC | MP033 30.6.2015 | - | - | +++ | 1:1 | - | - |
| | | | | | 4.7:1 | | |
| | | | | | 47:1 | | |
| 2 | 2016 | + | - | +++ | 2.55:1 | + | + |
| | | | | | 10:1 | - | - |
| 3 | 4007 | missing | | +++ | 2.55:1 | - | - |
| | | | | | 10:1 | +++ | +++ |
| **4** | 2027 (Fresh) | - | - | +++ | 4.3:1 | - | - |
| | | | | | 8.7:1 | +++ | ++ |
| | | | | | 17.4:1 | - | + |
| | 2027 | - | - | +++ | 1 | + | - |
| | | | | | 2 | ++ | - |
| | | | | | 3 | - | - |
| **5** | 2011 | - | - | +++ | 4.3:1 | - | - |
| | | | | | 8.7:1 | - | - |
| | | | | | 17.4:1 | + | +++ |
| 6 | 2001 | + | - | +++ | 4.3:1 | - | ++ |
| | | | | | 8.7:1 | +++ | ++ |
| | | | | | 17.4:1 | +++ | Missing |
| 7 | 2010 | - | - | +++ | 4.3:1 | - | ++ |
| | | | | | 8.7:1 | - | ++ |
| | | | | | 17.4:1 | + | |
| | 2010 repeat | + | - | +++ | 1 | + | - |
| | | | | | 2 | - | - |
| | | | | | 3 | - | - |
| 8 | 2018 | + | - | +++ | 4.3:1 | - | - |
| | | | | | 8.7:1 | - | +++ |
| | | | | | 17.4:1 | ++ | + |
| 9 | 2021 | + | - | +++ | 1 | +++ | + |
| | | | | | 2 | +++ | + |
| | | | | | 3 | - | - |
| 10 | 2025 | missing | + | +++ | 1 | missing | - |
| | | | | | 2 | | ++ |
| 11 | 4001 | +++ | - | +++ | 1 | +++ | + |
| | | | | | 2 | +++ | ++ |
| | | | | | 3 | +++ | ++ |
| 12 | 2031 | - | - | +++ | 1 | - | - |
| | | | | | 2 | +++ | +++ |
| | | | | | 3 | + | + |

**Figure S10:** A summary table of the responses from multiple repeats using human patient samples. The first sample (patient MP033) is the response detected when the donor is known to be DR4 negative, so no response is recorded. The control columns refer to the addition of soluble peptides or INFRANIX as a positive control for activation. No response (-) is determined as a response which is less than three folds above the no peptide background (or no MHC for the LbL samples). (+, ++ and +++ refer to responses of varying strength from low to very high). The cell to LbL ratios differed between different donors and different batches and hence the information there should be taken for qualitative purposes only.
